# Downregulation of GPR183 on Infection Restricts the Early Infection and Intracellular Survival of Mycobacterium tuberculosis in Macrophage

**DOI:** 10.1101/832592

**Authors:** Jun Tang, Ya’nan Shi, Lingjun Zhan, Chuan Qin

**Affiliations:** Institute of Laboratory Animal Sciences, Chinese Academy of Medical Sciences (CAMS), Comparative Medicine Center, Peking Union Medical College (PUMC), Beijing, China; NHC Key Laboratory of Human Disease Comparative Medicine, China; Beijing Key Laboratory for Animal Models of Emerging and Reemerging Infectious, China; Beijing Engineering Research Center for Experimental Animal Models of Human Critical Diseases, China; Tuberculosis Center, Chinese Academy of Medical Sciences (CAMS), Beijing, China

**Author notes:** Correspondence should be addressed to Lingjun Zhan and Chuan Qin.

**Keywords:** GPR183, EBI2, Mycobacterium tuberculosis, RAW264.7, BMDM

## Abstract

GPR183/EBI2 is a key chemotactic receptor for the positioning of B cells in lymphoid organs, and also for the migration of T cells and other immune cells. Here, we demonstrate that the downregulation of GPR183 in macrophage induced during Mtb infection restrains the bacterial early infection and intracellular survival. Overexpression of GPR183 or stimulation with its natural ligand favors Mtb survival in macrophage, while treatment with its antagonist represses both Mtb early infection and intracellular survival. With mutational analysis, we find that substitution of Asp-73, Arg-83, Tyr-112, Tyr-256 abolished the promotive effect of GPR183 on Mtb early infection and survival in macrophage. In conclusion, we demonstrated that beside the known role of chemotaxis receptor, GPR183 also functions directly in the interaction between macrophage and Mtb in a cell-autonomous way.

## Introduction

G-protein coupled receptors (GPCRs) widely participate in human physiology and are the most common therapeutic targets of approved chemical drugs. In the host-microbial interactions, they could also be the key mediators. N-acyl-amides from commensal bacteria could mimic human endogenous ligands and interact with gastrointestinal ‘lipid-like’ GPCRs, such as GPR119 and S1PR4, to alter the host physiology like blood glucose et al[1]. To the pathogenic microbes, GPCRs could act as chemotaxis receptors of immune cells, pattern recognition receptor (PRR) to bacterial components, and microbicidal mediators [2, 3]

Recently, several GPCRs have been proved to play roles in mediating signal transduction during mycobacterium infection. Some are manipulated by Mtb to facilitate the niche formation of the pathogen. Mtb infected monocytes promote increased MMP-1 and reduced TIMP-1 secretion of pulmonary epithelial cells under the synergy of TNFα and some GPCR, contributing to the tissue destruction and cavitation[4]. Virulent Mtb infection could enforce the feedback activation of the antilipolytic GPCR GPR109A, leading to accumulation of lipid bodies in foamy macrophages[5]. Others are necessary in the host immune responses to Mtb infection. Chemokine receptor CCR5 recognizes mycobacterial Hsp70 and mediates the following stimulation of human immature dendritic cell[6]. Mycobacterial secreted protein CFP-10 could trigger chemotaxis and reactive oxygen species production in neutrophil in a GPCR dependent manner[7]. In a host-targeted chemical screen, the identified inhibitors that could restrict mycobacterial growth in macrophage includes a group of GPCR modulators, suggesting more possible roles of GPCR in the host-pathogen interaction[8].

GPR183/EBI2 is found to act as a chemotactic receptor for B cells, T cells, splenic dendritic cells, macrophages and astrocytes[9–11]. It was de-orphanized in 2011 on the verification of 7α,25-dihydroxycholesterol (7α,25-OHC) to be its main natural ligand[12, 13]. Two synthetic enzymes CH25H and CYP7B1 and degrading enzyme HSD3B7 of 7α, 25-OHC collaboratively control the ligand concentration in lymphoid tissues and consequently guide B cell positioning and humoral immune responses[14]. 7α,25-OHC-GPR183 signaling is also critical to group 3 innate lymphoid cells (ILC3s) migration and lymphoid tissue formation by ILC3s in colon, and to the protective immunity to enteric bacterial infection[15, 16]. In innate immune cells such as macrophages, oxysterol could also direct the migration of cells through GPR183[11]. In 2010, a research identified 275 genes including GPR183 regulating Mtb load in human macrophages by a genome-wide siRNA screen[17], while no further investigation on the role of GPR183 in tuberculosis has been reported since then. In previous study, we found the downregulation of Gpr183 mRNA in the whole blood of Mtb infected mice. Here we further describe the functional role of GPR183 in macrophage during Mtb infection.

## Materials and methods

### Cell culture, infection and intracellular bacilli load determination

Plated RAW264.7 cells were infected with Mtb H37Rv at a multiplicity of infection of 10 or 25 during 3 h of infection followed by 40μg/mL gentamycin treatment for 1 h to remove the remaining extracellular bacilli. Mouse BMDMs from C57BL/6 or *Ifnar1^−/−^* mice [18] were obtained from bone marrow and allowed to differentiate with 10ng/mL mM-CSF for 7 days. For cfu determination, infected cells were lysed with 0.05% aseptic SDS and plated on Lowenstein-Jensen (LJ) slants at serial dilutions. The cfu was enumerated following incubation at 37°C for 21-28 days. For time-to-detection determination, serial dilutions of the above cell lysis were added into MGIT 960 tubes and monitored by BACTEC MGIT 960 system as previously described[18].

### Western blot and antibodies

Cell lysates were generated at defined time-points with RIPA lysis buffer containing protease inhibitor cocktail (Thermo Scientific). GPR183 rabbit pAb (Abcam ab229527), GAPDH mouse mAb (Proteintech HRP-60004), CH25H rabbit pAb (Abcam ab133933), CYP7B1 rabbit pAb (Proteintech 24889-1-AP) were used for probing and the blots were recorded with a CCD imaging workstation (ChemiDoc, Bio-rad).

### RNAi-mediated silencing

siRNAs were designed and synthesized by RiboBio (Guangzhou, China). A mix pool of siRNAs was transfected into RAW264.7 with HiPerFect Reagent (Qiagen) and the knockdown efficiency determination or the following assays were carried out 48 h later. SiR NC, 5’-UUCUCCGAACGUGUCACGU-3’. Si-m-Gpr183-1, 5’-GGCUAACAAUUUCACUACC-3’. Si-m-Gpr183-2, 5’-CAACCACUCUCUAUUCAAU-3’. Si-m-Gpr183-3, 5’-AGACAUUCCUUCCAGAUCU-3’

### Plasmids construction and overexpression

The Gpr183 expression plasmid (Origene MR205447) was used as the wildtype control and underwent site-directed mutagenesis, including D73A, R83A, Y108A, Y112A, Y256A and D73A/R83A. All mutations were verified using DNA sequencing. The plasmids were transfected into RAW264.7 with Attractene Reagent (Qiagen) and the overexpression efficiency determination or the following assays were also carried out 48 h later.

### Real time PCR

Cells were lysed with TRIzol Reagent and the total RNA was isolated according to the instructions of the manufacturer. The cDNA was obtained using RevertAid RT Reverse Transcription Kit (Thermo Scientific) and used as the template in the following Real time PCR assay. Gpr183 primers, 5’-AATCAACTCAACCACTCT-3’ (Forward), 5’-ATCACCTATCCTCCAATC-3’ (Reverse). Gapdh primers, 5’-GAGTCTACTGGTGTCTTC-3’ (Forward), 5’-AATCTTGAGTGAGTTGTC-3’ (Reverse). Ch25h primers, 5’-CATCACTCTCAGTTTAAC-3’ (Forward), 5’-TTCTTCTTCTGGATCAAG-3’ (Reverse). Cyp7b1 primers, 5’-AATAGGAGCACATCATCT-3’ (Forward), 5’-GCCGAAGAATATAATACATTG-3’ (Reverse). Hsd3b7 primers, 5’-AGAGACTTCTACTACCAG-3’ (Forward), 5’-AAGGTGACTTATCATAGC-3’ (Reverse). The relative expression levels were determined using StepOne plus (Applied Biosystems) and the ΔΔCt method.

### Statistical Analysis

Experiments were repeated two or three times. One-way ANOVA and Student’s two-nailed *t*-test were used for statistical analysis.

## Results

### Mtb infection decreases GPR183 expression in vivo and in vitro

We have previously performed RNA sequencing to the whole blood from different groups of C57BL/6 mice, including healthy control, infected mice with Mtb H37Rv for six weeks, and treated mice receiving isoniazid for four weeks after infection. It is noticed that the expression of Gpr183, a G protein coupled receptor, is significantly down-regulated in Mtb infected group and stay low in treated group to a less extent (data not shown). Then the expression changes of Gpr183 mRNA was confirmed with qPCR (**Fig. 1A**). As the RNA profile of whole blood could be largely ascribed to monocytes and granulocytes, we further investigate the functional role of GPR183 in macrophage, the main target cell during Mtb infection. RAW264.7 cell was infected with H37Rv to provide an in vitro model to study macrophage infection.

**Figure 1.**
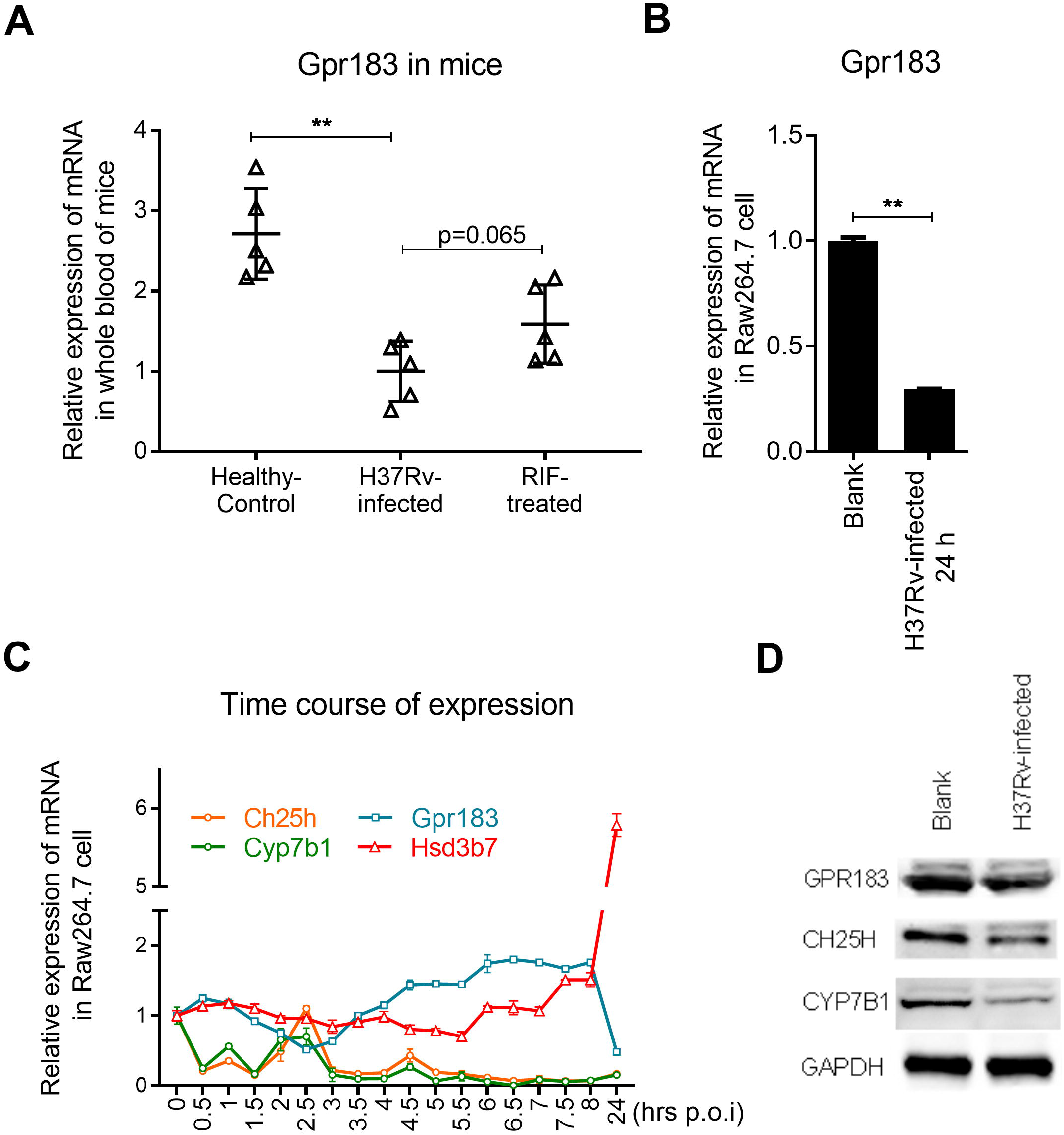
Mtb infection decreases GPR183 expression in vivo and in vitro. (A) Gpr183 mRNA level in whole blood of C57BL/6 mice (n=5), quantitated by real time PCR. (B) Expression of Gpr183 mRNA and protein (D) was downregulated in Mtb infected RAW264.7 24 h postinfection. (C) The mRNA changes of Gpr183 and its ligand related genes in the early infection phase were monitored in RAW264.7. (D) Protein levels of GPR183, CH25H and CYP7B1 were detected by immunoblotting in RAW264.7 24 h postinfection. **p<0.01.

In RAW264.7 cells infected with H37Rv for 24 h, the expression of Gpr183 is also significantly reduced (**Fig. 1B**). This is consistent with previous work which suggested the down-regulation of Gpr183 mRNA in murine macrophages on immune challenges such as LPS [19]. It is also reported that the regulation of Gpr183 mRNA on LPS stimulation is fast[11, 20]. Therefore, we examined the detailed time course of its expression with 30 min intervals, as well as the expression of the three enzymes closely related to 7α,25-OHC (**Fig. 1C**). There is a plunge of Gpr183 mRNA at 2.5 h post infection, followed with a slow climb, turning into upregulation within 8 h post infection, which is different from the response to LPS stimulation[11, 20]. In correspondence with that, there is a peak of cholesterol hydroxylases CH25H and CYP7B1 mRNA at 2.5 h post infection, and notable decrease at 24 h poi, suggesting the 7α,25-OHC-GPR183 signaling may under collaborative regulation on Mtb infection, although CH25H and CYP7B1 decrease earlier and more drastically than GPR183 at both of mRNA and protein levels (**Fig. 1D**). The dehydrogenase HSD3B7 mRNA changes a little within 8 h post infection, but significantly increased at 24 h poi. When performing its role in T cell and B cell chemotaxis, GPR183 receives 7α,25-OHC mediated downregulation, which is important to the correct positioning of T cell and B cell[14, 21]. Then these transcriptional expression changes may suggest a functional role of GPR183 during Mtb infection.

### Downregulation of GPR183 restrains the early infection and the survival of Mtb in macrophage

Then we overexpressed GPR183 with plasmid in RAW264.7 cell and examined its effect to the early infection and the survival of Mtb in host cell. First, the colony forming unit (CFU) of Mtb in RAW264.7 cells was determined at 4 h after Mtb being added to the cultured cell. This early infection process includes phagocytosis of Mtb and partial restrained stage of Mtb in phagosome. In this process, overexpression of GPR183 increased the CFU by about 50% over the vector transfected group (**Fig. 2A**). Besides, the CFU of Mtb in cells was determined at 48 h after Mtb being added into the cell culture. The transfection of plasmid was performed after Mtb infection, so that the effect of overexpression on the survival of Mtb in macrophage could be detected. In this process, overexpression of GPR183 also increased the CFU over the vector group (**Fig. 2B**). Beside the CFU counting on plates, the cell lysates were also cultured using MGIT 960 system, and Time to Detection (TTD) of the Mtb in cell scored by the system was detected and compared among groups. Briefly, shorter TTD indicates higher bacterial load (**Fig. 2A, B** right panels). In accordance with the CFU results, TTD of GPR183 overexpression groups were significantly shorter than the vector groups.

**Figure 2.**
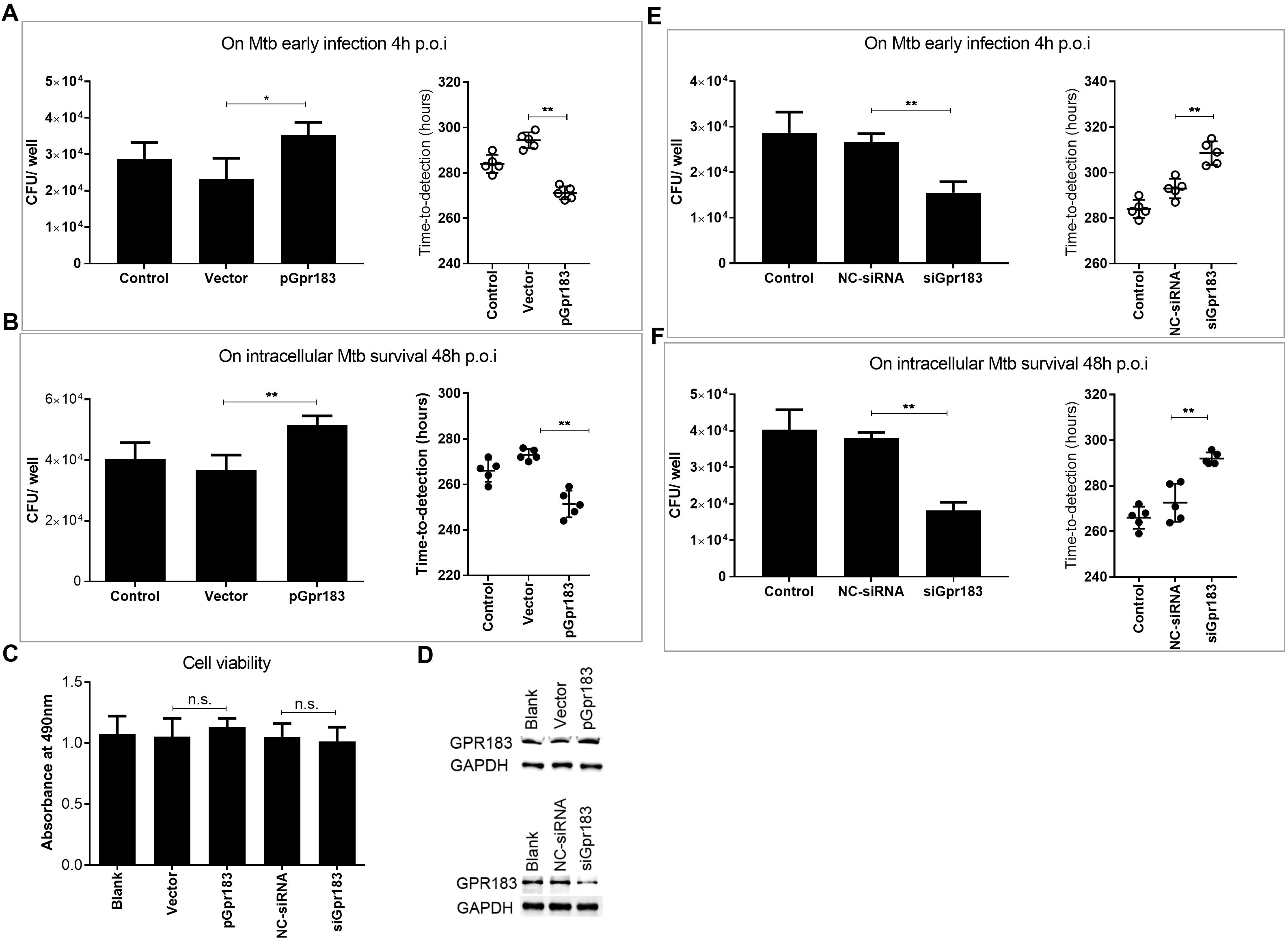
Mtb-induced downregulation of Gpr183 represses bacterial early infection and survival in macrophage. Forced expression of Gpr183 enhanced Mtb early infection (4 h poi) (A) and survival (48 h poi) (B) in RAW264.7. (C) The cell viability at 48 h after transfection showed no significant difference determined by MTS assay. (n.s. p>0.05). (D) Transfection efficiency of overexpression plasmid and siRNAs in RAW264.7 was confirmed by immunoblotting. Blots are representative of three independent experiments. Knockdown of Gpr183 repressed Mtb early infection (4 h poi) (E) and survival (48 h poi) (F) in RAW264.7. CFU determination is presented in the left panels, and time-to-detection (TTD) scored by MGIT960 system is presented in the right panels. (n=5). *p<0.05, **p<0.01.

The knockdown of GPR183 was performed with a pool of three siRNA. Neither overexpression nor knockdown affected the cell viability of cultured cells (**Figure 2C, D**). The knockdown effect to the infection and survival of Mtb in host cell was also examined as in the overexpression assay. In both of the early infection and the following survival process of Mtb in R264.7 cells, knockdown of GPR183 decreased the CFU of intracellular Mtb, compared with the Negative Control (NC) siRNA groups (**Fig. 2E and 2F**). The GPR183 knockdown groups also had longer TTD than NC groups. These data indicated a promotive role of GPR183 in the early infection and the survival of Mtb in macrophage.

### The antagonist NIBR189 could suppress the early infection and the survival of Mtb in macrophage

As the potent natural ligand of GPR183 has been identified and a specific antagonist, NIBR189, has been developed[22], we further tested the effect of these small molecules on the early infection and the survival of Mtb in macrophage. Firstly, the cell viability under a wide range of concentrations of these molecules was tested and was found to have little difference (**Figure 3A**). At the presence of 25nM NIBR189, the CFU of intracellular Mtb at 4 h after being added to RAW264.7 cells decreased about 40% over infection control group (**Fig. 3B**). Then 25nM NIBR189 was added into Mtb-infected RAW264.7 cells to determine its effect on the CFU at 24 h, 48 h and 72 h post infection. It was observed that NIBR189 also reduced the intracellular Mtb at the stages after phagocytosis (**Fig. 3C**). The results indicated a similar effect of NIBR189 treatment with that of GPR183 knockdown, which confirmed the function of GPR183 in the early infection and the survival of Mtb in macrophage and provided a potential candidate for host-targeted treatment of tuberculosis.

**Figure 3.**
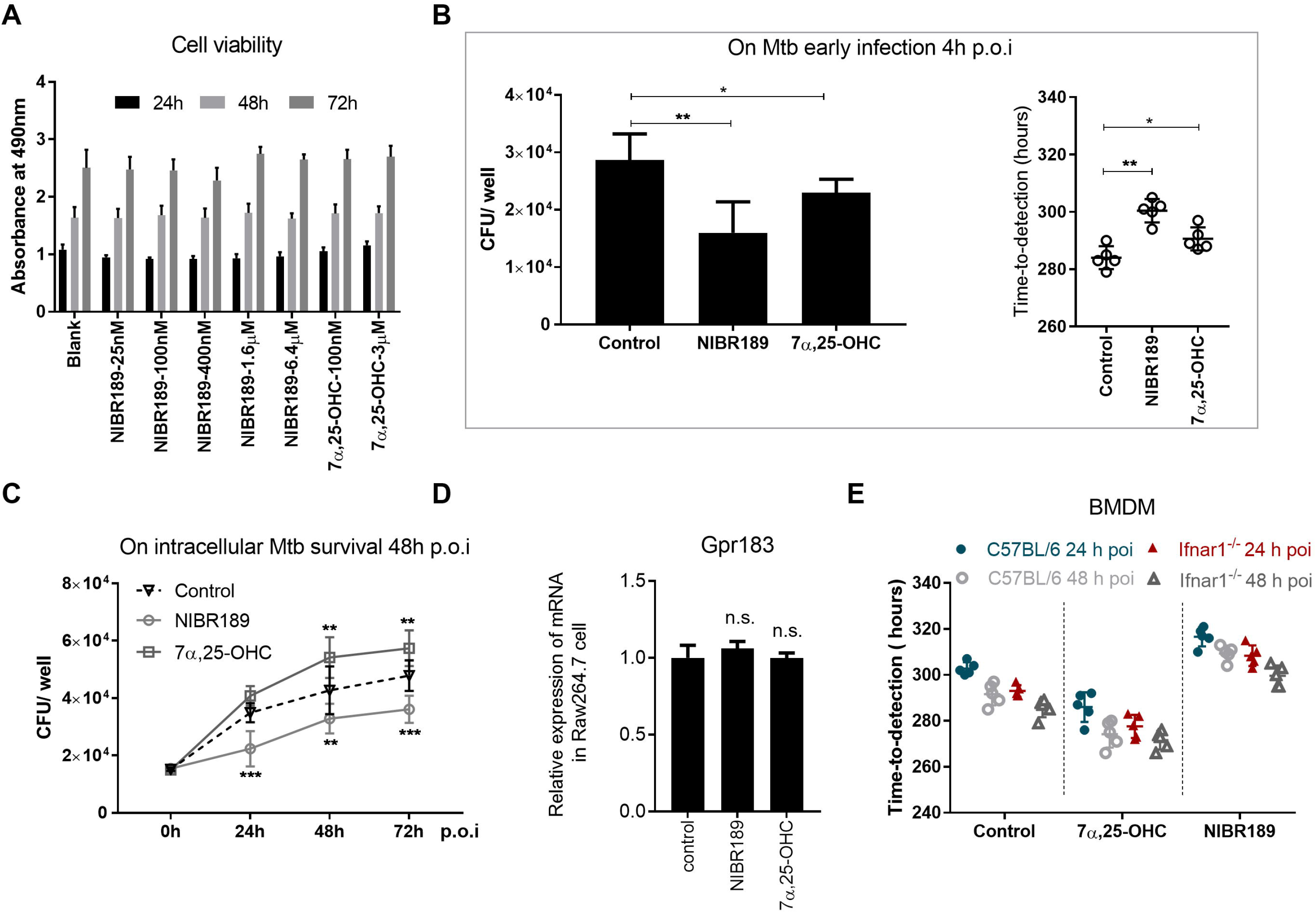
Role of chemicals targeting GPR183 in the early infection and the survival of Mtb in macrophages. (A) The cell viability under the treatment of NIBR189 and 7α,25-OHC at different time points changes little, determined by MTS assay. (B) Bacterial early infection (4 h poi) is compromised under the treatment of NIBR189 and 7α,25-OHC in RAW264.7. (CFU in left panel, TTD in right panel) (C) The effect of NIBR189 and 7α,25-OHC on intracellular Mtb survival is compared using cfu at different time points after infection in RAW264.7. (D) NIBR189 and 7α,25-OHC exert little modulation on Gpr183 mRNA expression. n.s. p>0.05. (E) BMDMs from C57BL/6 and *Ifnar1^−/−^* mice were infected with Mtb for 24 and 48 h, under the treatment of 25nM NIBR189 or 100nM 7α,25-OHC. The time-to-detection scored by MGIT960 system is compared between groups. * p<0.05, **p<0.01, ***p<0.001.

The effect of the natural ligand 7α,25-OHC was also tested as in the antagonist assay. Surprisingly, 100nM 7α,25-OHC also reduced intracellular Mtb at the early infection process like NIBR189, while increased intracellular CFU at 24 h, 48 h and 72 h post infection (**Fig. 3B and 3C**). The different effect of 7α, 25-OHC in the early infection process and the survival process suggests that its roles in the two stages may involve different mechanisms. Although the mRNA level of Gpr183 was not affected on the treatment of NIBR189 or 7α,25-OHC (**Figure 3D**), the internalization of GPR183 after binding with 7α,25-OHC, which is the main mechanism of its desensitization, could result in the downregulation of GPR183 at the cell surface[13]. The reduced availability of surface GPR183 may partially explain the early suppression of 7α,25-OHC to the bacilli infection.

As it is reported that GPR183 could negatively regulate type I interferons in plasmacytoid and myeloid dendritic cells[23], we tested the effect of NIBR189 and 7α,25-OHC on the survival of Mtb in bone marrow - derived macrophages (BMDMs) from C57BL/6 mice and *Ifnarf^−/−^* mice. Similar to the results in RAW264.7 cells, in BMDMs both from C57BL/6 mice and *Ifnar1^−/−^* mice, 25nM NIBR189 reduced the intracellular Mtb, while 100nM 7α,25-OHC increased the bacilli load at 24 h and 48 h post infection, which is indicated by TTD (**Fig. 3E**). Therefore, the roles of GPR183 agonist and antagonist during Mtb infection are performed through type I IFN - independent manners in BMDMs.

### Introduction of key residue mutations abolished the promotive effect of GPR183 on Mtb early infection and survival in macrophage

To further investigate the molecular basis of the promotive role of GPR183 in Mtb infection, we performed mutational analysis to identify the important residues. Arg-83, Tyr-108, Tyr-112, Tyr-256, which are important for agonist binding in human GPR183, and Asp-73, which is essential for the activation mechanism of human GPR183 but not for 7α,25-OHC binding [24], were separately substituted with Ala to construct receptor mutants. A double mutant D73A/R83A was also generated to examine the effect in combination (**Fig. 4A**). The intracellular bacterial load at the early infection phase and the survival phase in RAW264.7 transfected with these constructs, wild type receptor or vector was determined through MGIT 960 system. On Ala substitution of Asp-73, Arg-83, Tyr-112, Tyr-256 or D73/R83, the effect of overexpression of GPR183 on the intracellular Mtb was abolished (**Fig. 4B and 4C**). These results further suggest that the GPR183 activation, not just ligand binding, may play important roles during Mtb infection.

**Figure 4.**
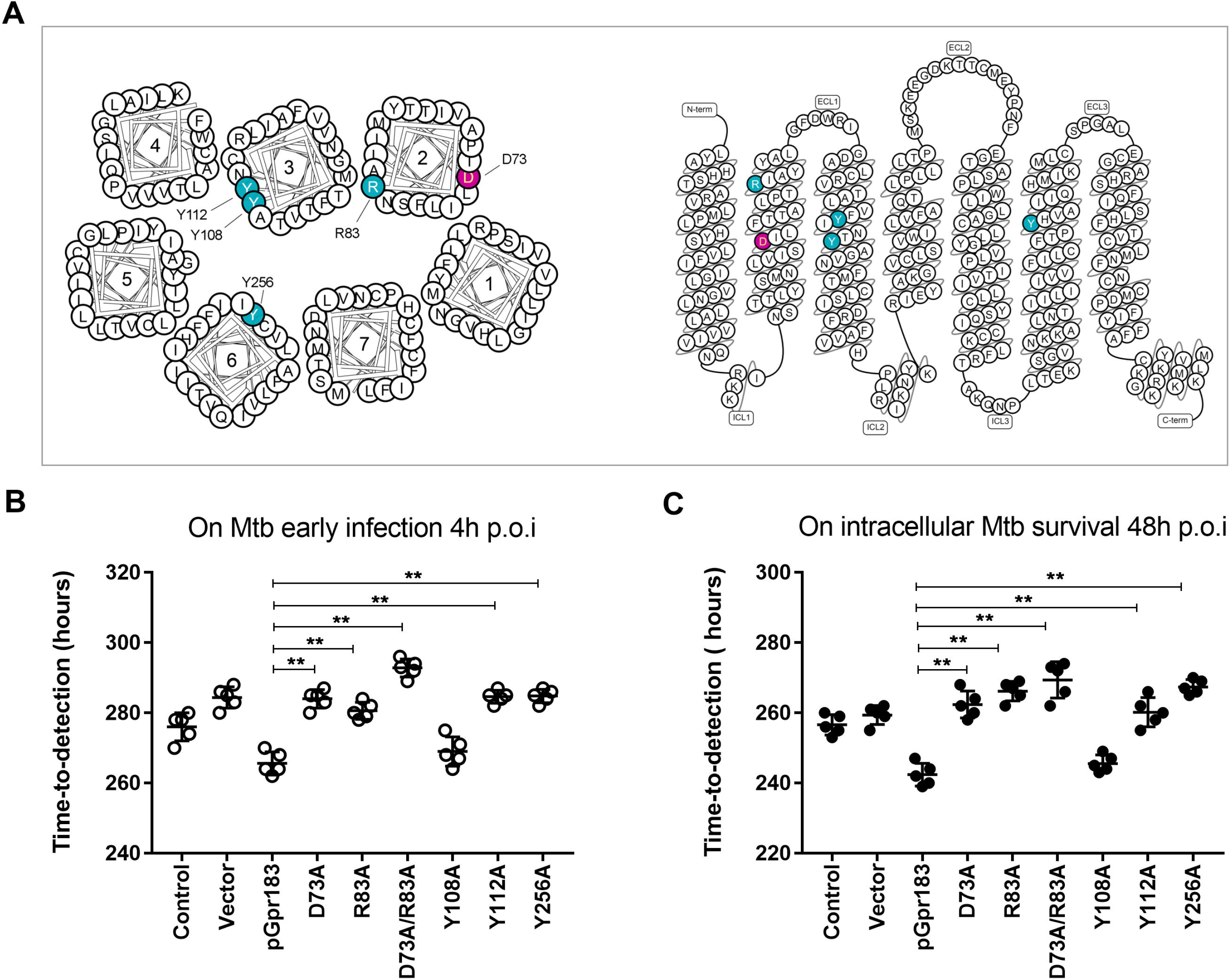
Mutational analysis on the role of GPR183 during Mtb infection. (A) The helical wheel (left panel) and snake-diagram (right panel) of the mouse GPR183 are modified from GPCRdb. Mutated residues are indicated in blue (7α,25-OHC binding points in human homolog) or in red (effect on receptor activation in human homolog). The effect of mutants’ overexpression on Mtb early infection (4 h poi) (B) and intracellular Mtb survival (48 h poi) (C) is compared with that of wildtype Gpr183, using TTD scored by MGTI960 system. Shorter TTD indicates higher bacterial load. **p<0.01.

## Discussion

In this study we demonstrated that GPR183 could play a promotive role in macrophage during Mtb infection process and inhibition of GPR183 activity could repress the early infection and intracellular survival of Mtb. On Mtb infection, Gpr183 received downregulation in mouse whole blood and macrophage cell line RAW264.7, so were its ligand synthetic enzymes Ch25h and Cyp7b1. These expression patterns were different from that in macrophage under LPS stimulation[25, 26]. This downregulation of 7α,25-OHC-GPR183 signaling in macrophage may act as a beneficial response for host immunity to control Mtb infection, considering the key role of macrophage in tuberculosis. Even so, further inhibition of GPR183 activity could be considered as a host-targeted intervention to control Mtb infection.

As a natural ligand of GPR183, 7α,25-OHC delivers delicate signals to various cells depending on its concentration gradient. In this study, 7α,25-OHC also reduced intracellular Mtb like antagonist NIBR189 at the early infection process which mainly includes phagocytosis and restrained stage of Mtb in phagosome. This interesting observation is not due to decreased cell viability. After binding with its ligand, GPR183 is reported to get internalized into cells, leading to its downregulation from cell surface[13, 27], which is a major mechanism of the desensitization of GPR183. According to our knockdown and overexpression assays, the expressing level of GPR183 is important to the early infection process of Mtb. In addition, the receptor internalization could happen fast[27], therefore it may partially explain the effect of 7α,25-OHC on the early infection.

Although the function of GPR183 in B cell has been intensively investigated[13], the role and the mechanism of GPR183 in macrophage has not been well elucidated[11], especially in the interaction between macrophage and bacteria. Beside the role of chemotaxis receptor[20], here we demonstrated that GPR183 also functions directly in the interaction between macrophage and Mtb in a cell-autonomous way. Using *Ifnar1^−/−^* BMDMs we excluded type I interferons as the downstream effectors of GPR183, and the downstream mechanism still warrants further investigation.

## Acknowledgments

This work was supported by National Science and Technology Major Project for Infectious Diseases Control and Prevention [grant number 2017ZX10304402-001-016]; National Natural Science Foundation of China [grant number 31600742]; PUMC Basic Research Grant [2017PT31010]; and CAMS Initiative for Innovative Medicine [grant numbers 2016-I2M-1-013 and 2016-I2M-2-006].

